# Perinatal exposure to fluoxetine and maternal adversity affect myelin-related gene expression and epigenetic regulation in the corticolimbic circuit of juvenile rats

**DOI:** 10.1101/2020.07.26.221648

**Authors:** Anouschka S. Ramsteijn, Rikst Nynke Verkaik-Schakel, Danielle J. Houwing, Torsten Plösch, Jocelien D.A. Olivier

## Abstract

Many pregnant women experience symptoms of depression, and are often treated with selective serotonin reuptake inhibitor (SSRI) antidepressants, such as fluoxetine. *In utero* exposure to SSRIs and maternal depressive symptoms is associated with sex-specific effects on the brain and behavior. However, knowledge about the neurobiological mechanisms underlying these sex differences is limited. In addition, most animal research into developmental SSRI exposure neglects the influence of maternal adversity. Therefore, we used a rat model relevant to depression to investigate the molecular effects of perinatal fluoxetine exposure in male and female juvenile offspring. We performed RNA sequencing and targeted DNA methylation analyses on the prefrontal cortex and basolateral amygdala; key regions of the corticolimbic circuit. Perinatal fluoxetine enhanced myelin-related gene expression in the prefrontal cortex, while inhibiting it in the basolateral amygdala. SSRI exposure and maternal adversity interacted to affect expression of genes such as myelin−associated glycoprotein (*Mag*) and myelin basic protein (*Mbp*). We speculate that altered myelination reflects altered brain maturation. In addition, these effects are stronger in males than in females, resembling known behavioral outcomes. Finally, *Mag* and *Mbp* expression correlated with DNA methylation, highlighting epigenetic regulation as a potential mechanism for developmental fluoxetine-induced changes in myelination.

## Introduction

The use of selective serotonin reuptake inhibitor (SSRI) antidepressants during pregnancy has greatly increased^1–4^. Recent estimates suggest that every year, in Europe and the US alone, hundreds of thousands of babies are born that have been exposed to SSRI medication^5–8^. SSRIs cross the placental barrier^9^ and target the fetal serotonin transporter (SERT). This potentially affects brain circuit formation, as serotonin serves as a neurotrophic factor and influences neurogenesis, cell migration, axon guidance, dendritogenesis and synaptogenesis during development^10^. Some studies find no long-term associations between *in utero* SSRI exposure and cognitive and behavioral outcomes^11–13^. However, others have reported higher anxiety^14^, lower motor-, social-emotional- and adaptive behavior^15^ and a higher risk of developing mental and behavioral disorders^16^ after prenatal SSRI exposure. Some of these associations are modulated by offspring sex^17,18^. For example, an increased risk for autism spectrum disorder was only found for boys^19^. A limitation of this field is the confounding factor of maternal depression^16^, which has been associated with neurodevelopmental outcomes resembling those of SSRI exposure^20^. In fact, one study found that a higher risk of autism spectrum disorder after SSRI exposure during gestation did not hold up after controlling for maternal psychiatric illness, suggesting it was the underlying maternal disorder that is responsible for the effect^21^. In addition, prenatal maternal depression is related to neurobehavioral function in neonates^22^, motor development and language scores in infants^23^, and executive functioning capabilities in 6-year-olds^12^. The relative contribution of maternal SSRI use versus the underlying depression is often unclear.

Neuroimaging techniques are used to investigate the effects of *in utero* exposure to SSRIs and maternal depressed symptoms on brain structure and function in humans. Prenatal SSRI exposure is associated with altered white and gray matter architecture^24^ and connectivity^25–27^ in babies. Even though the SSRI-exposed group is usually compared to a control group exposed to unmedicated depressive symptoms of the mother^25–27^, it remains possible that the intensity or nature of the underlying depressive symptoms differs between these groups and partially mediates the observed effects^28^. It is known that the severity of antenatal depressive symptoms correlates to connectivity in brain networks involved in emotion regulation in infants^29^. Several studies report sex differences^30^, with girls showing stronger associations of prenatal exposure to depressive symptoms with functional connectivity in emotion-regulation networks^31^, right amygdala volume^32^, and cortical thickness^33^ than boys. Overall, prenatal exposure to SSRIs and maternal depression affect the developing brain in overlapping-but potentially also in distinct brain regions^34^, with a key role for corticolimbic structures such as the prefrontal cortex and the amygdala^25,35,36^.

Rodent experiments offer the ability to causally investigate the separate and combined effects of SSRI administration and aspects of maternal depressed mood. Neurodevelopmental patterns^37^, including serotonin system-specific ones^38–40^, are remarkably conserved between rodents and humans, with the first postnatal weeks in rodents corresponding to the third trimester of pregnancy in humans^41^. In a meta-analysis of rodent behavioral outcomes, we showed that perinatal SSRI exposure is associated with reduced activity and exploration behavior, a more passive stress-coping style, and less efficient sensory processing^42^. The neurobiological correlates of these behavioral changes likely include the serotonergic system^43^. However, the effects of perinatal SSRI exposure on the brain are more widespread, with roles for the prefrontal cortex^44,45^, limbic structures^46^, and the dorsal raphe nucleus^45,46^. Rodent models of maternal depressive-like symptoms are able to recapitulate basic neurobiological changes seen in women with peripartum depression and their children^47^. *In utero* exposure to maternal stress alters behavioral outcomes in rodents, as was shown for anxiety-like behavior and learning and memory performance^48^. Sex-specificity is often seen for neurodevelopmental outcomes after both perinatal SSRI exposure^49–51^ and prenatal stress^49,52^. Interestingly, perinatal SSRI exposure might reverse some of the neurodevelopmental effects of maternal (pre)gestational stress, such as altered brain serotonin levels^49,53^ and stress-coping behavior^54^.

In contrast to human studies, rodent studies enable investigation of molecular changes during fundamental neurodevelopmental periods that may trigger these long-term effects. Perinatal SSRI exposure alters serotonin system-related gene expression in mice^55^ and rats^56^. Reinforcing the idea that sex is a modulating factor, early postnatal FLX exposure and prenatal stress *decreased* brain-derived neurotrophic factor (*Bdnf*) exon 4 mRNA levels in adult males^57^, while prenatal stress *increased Bdnf* exon 4 expression in females^58^. Transcriptome-wide gene expression has been studied with microarrays. Whereas *prenatal* SSRI exposure and/or stress showed no effects in the amygdala, hippocampus or hypothalamus^59^, early *postnatal* SSRI exposure altered transcriptomic status of the hippocampus^60^. Developmental screening of gene expression revealed that the hippocampus showed the largest paroxetine-induced differences in the first 2 postnatal weeks with changes in neurogenesis, synaptic structure and function, and energy metabolism pathways^61^. The amygdala showed the largest differences around PND21^61^, suggesting that the transcriptomic response to SSRIs depends on brain region and developmental time window. Despite evidence for sex-specific effects, these transcriptome-wide analyses have only been done using male animals^59–61^, and only one study considered prenatal stress^59^. In addition, whereas the hippocampus is a widely studied brain area, other regions in the corticolimbic circuitry are underrepresented in research.

To address these gaps in the literature, we aimed to investigate transcriptomic alterations in the corticolimbic circuitry of male and female juvenile rats exposed to maternal adversity and/or perinatal SSRIs using RNA sequencing. To this end, we used a genetic rat model of maternal vulnerability (MV). MV females show a depressive-like phenotype in adulthood after being exposed to early life stress (sMV). This is reflected in anhedonic behavior and lower central neural growth factor gene expression in sMV-than in control (cMV) females^62^. We treated sMV and cMV dams daily with fluoxetine (FLX), a commonly used SSRI, or vehicle (Veh) from the start of gestation until weaning at postnatal day (PND)21 of the pups. We have previously reported sex-specific effects of developmental FLX exposure on social behaviors in this model, with males being more vulnerable^63^. To investigate the molecular mechanisms underlying these findings, we collected brains from male and female offspring at PND21 for RNA sequencing of micropunched tissue from the medial prefrontal cortex (PFC) and the basolateral amygdala (BLA). To examine a potential role for epigenetic regulation, we quantified DNA methylation levels of differentially expressed genes in the same tissue.

## Methods

### Experimental animals

Our rodent model of maternal vulnerability (MV) consists of heterozygous serotonin transporter knockout (SERT^+/−^) female rats (see below). Experimental animals were taken from our colony of serotonin transporter knockout (SERT^−/−^, Slc6a41^Hubr^) Wistar rats at the University of Groningen (Groningen, the Netherlands), and were derived from outcrossing SERT^+/−^ rats^64^. Animals were supplied with standard lab chow (RMH-B, AB Diets; Woerden, the Netherlands) and water *ad libitum* and were kept on a 12:12 h light-dark cycle (lights off at 11:00AM), with an ambient temperature of 21 ± 2 °C and humidity of 50 ± 5%. Cages were enriched with wooden stick for gnawing (10×2×2cm) and nesting material (Enviro-dri™, Shepherd Specialty Papers, Richland, MI, USA), and were cleaned weekly. Pregnant dams were housed individually in type III Makrolon cages (38.2×22.0×15.0cm) and stayed with their pups until postnatal day (PND)21. Pups were weaned at PND21 and group housed in same-sex cages of 3-5 animals in type IV (55.6×33.4×19.5cm) Makrolon cages. SERT genotyping was performed on ear punches, and was performed as described previously^65^. All experimental procedures were approved by the Institutional Animal Care and Use Committee of The University of Groningen and were conducted in agreement with the Law on Animal Experiments of The Netherlands.

### Maternal adversity and fluoxetine treatment

SERT^+/−^ female rats (MV) were exposed to adversity in the form of early life stress (sMV) to induce anhedonia^62,66^. The early life stress protocol was conducted as previously described^65^. In short, SERT^+/−^ animals were crossed (F0). Pregnant females were left undisturbed until delivery, which was defined as PND0. From PND2 to PND15, pups were either separated from the dam for 6 hours per day, or control handled 15 minutes per day. Treatment allocation was done in alternating fashion based on date of birth. The SERT^+/−^ female pups from these nests (F1) then matured to become sMV- and control (cMV) females (Figure 1).

**Figure 1:**
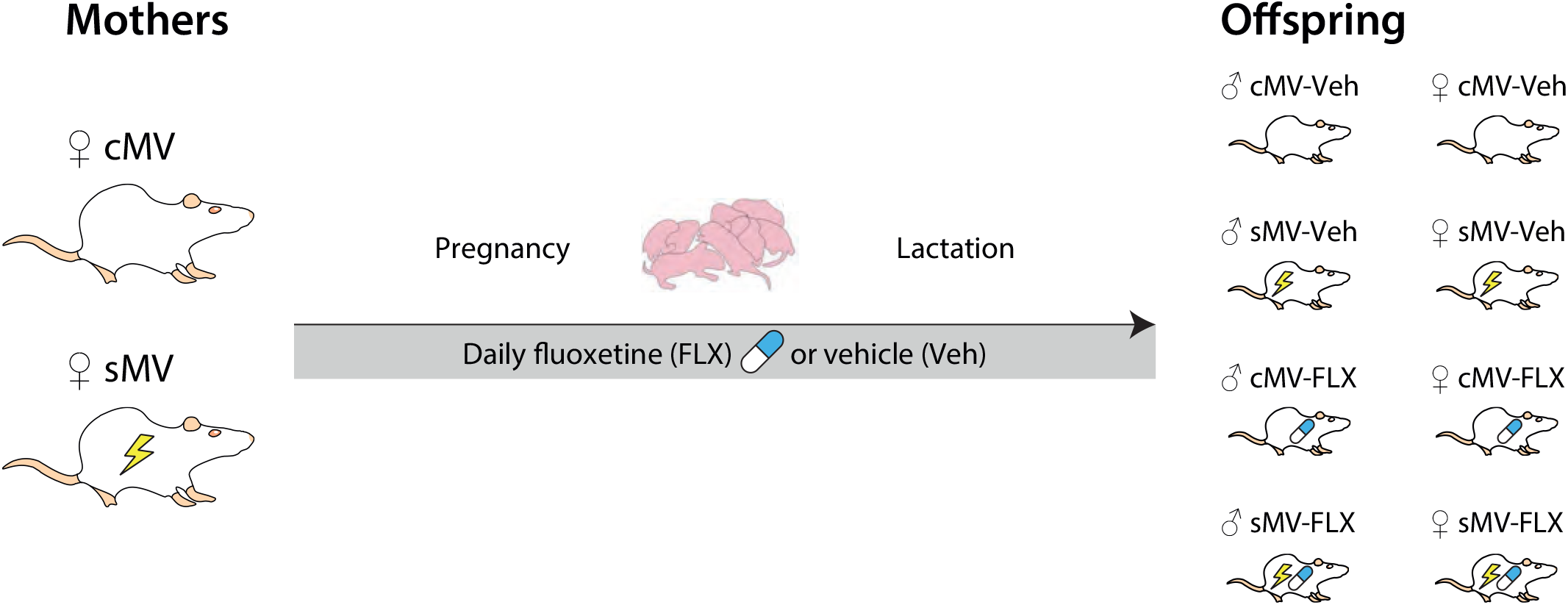
Overview of study design. Maternal vulnerability (MV) rats were either exposed to stress (sMV) or control handled (cMV) early in life. In adulthood, sMV and cMV females were bred. Throughout pregnancy and lactation, from gestational day (GD)1 until PND21, females received a daily oral injection of either 10 mg/kg fluoxetine (FLX) or methylcellulose (Veh). This resulted in 8 offspring groups: cMV-Veh males, cMV-Veh females, sMV-Veh males, sMV-Veh females, cMV-FLX males, cMV-FLX females, sMV-FLX males, and sMV-FLX females. Offspring brains were taken at PND21 for molecular analysis (N=5/group).

In adulthood, sMV and cMV females were mated with wildtype males. Females were mated when in estrus (checked with a disinfected impedance meter, model MK-11, Muromachi, Tokyo, Japan), between the age of 3 and 6 months. This was termed gestational day (GD)0. Males were removed after 24 hours, and females stayed isolated in a type III cage. Throughout pregnancy and lactation, from GD1 until PND21, the dams were weighed and received an oral gavage of either 10 mg/kg fluoxetine (FLX, Fluoxetine 20 PCH, Pharmachemie BV, the Netherlands) or vehicle (Veh) daily at 11:00AM (Figure 1). Treatment allocation was done semi-randomly; when possible, females from the same litter were put in different treatment groups. Methylcellulose (MC; Sigma Aldrich Chemie BV, Zwijndrecht, the Netherlands), the constituent of fluoxetine capsules, was used for vehicle injections. FLX (5 mg/mL) and MC (1%) solutions were prepared with autoclaved water. The gavage volume ranged from 0.9 mL to 2.0 mL, depending on body weight. Oral treatment was given by gently picking up the animal without restraint, and using flexible PVC feeding tubes (40 cm length, Vygon, Valkenswaard, the Netherlands) in order to minimize discomfort. It should be noted that our lab has been confronted with thus far unexplained mortality in about one fourth of MV females as a result of fluoxetine treatment, a phenomenon we have described elsewhere.^63^ None of the females included in the current study showed any signs of toxicity as a consequence of FLX, except for one. The pups from this sMV-FLX female who died at PND17 – when the pups are largely independent – were cross-fostered to a same-treated mother and were included in the study.

### Offspring brain collection and nucleic acid isolation

Pups from N=31 nests were used for this study (cMV-Veh N=9, sMV-Veh N=8, cMV-FLX N=8 and sMV-FLX N=6). Pups were checked and counted daily, and weighed weekly to track welfare and growth. Between PND14 and PND20, ear punches were taken for SERT genotyping, which was performed as described previously^65^. Only SERT^+/+^ (wildtype) offspring were used in this study. On PND21, approximately 24 hours after the last FLX or MC injection of the mother, N=8 males and N=8 female offspring per group (Figure 1) were killed by rapid decapitation. Brains were isolated, snap-frozen in isopentane (Acros Organics) on dry ice, and then stored at −80°C.

On a Leica CM3050 cryostat, 200 μM coronal brain sections were sliced. We made use of a stereotaxic atlas specifically developed for the PND21 rat brain to locate the brain areas of interest^67^. The medial prefrontal cortex (PFC) was collected from Bregma 2.8 mm to Bregma 1.8 mm (Supplementary Figure 1A). The basolateral amygdala (BLA) was collected bilaterally from Bregma −1.6 mm to Bregma −3.0 mm (Supplementary Figure 1B). A 2.0 mm punch (Harris Uni-Core) was used to collect these areas, which were then stored in 2.0 mL safe-lock tubes (Sarstedt) at −80 degrees Celsius.

To be able to examine gene expression and epigenetic regulation within the same samples, the AllPrep DNA/RNA Mini Kit (Qiagen) was used to simultaneously isolate DNA and RNA according to the manufacturer’s protocol. We started by adding 350 μL Buffer RLT Plus, 3.5 μL β-mercaptoethanol (Sigma-Aldrich) and 1.75 μL Reagent DX (Qiagen) per sample. Samples were lysed using a TissueLyser II (Qiagen) and 5 mm stainless steel beeds, 2 times for 2:00 at 30 Hz. After nucleic acid isolation, their concentration and purity were checked using a NanoDrop 2000 (Thermo Scientific). In addition, total RNA concentration and integrity were quantified using a RNA 6000 Nano Kit 2100 on a Bioanalyzer Instrument (Agilent Technologies, CA, USA).

### RNA sequencing

The RNA isolated from the PFC and BLA from N=5 male and N=5 female offspring per treatment group (N=80 samples in total) was selected for RNA sequencing. N=5 per group was expected to be sufficient to detect changes. Samples were selected based on RNA quantity and quality, and so that each group only included one animal per litter. The RNAseq, including library preparation, quality control and mapping of reads was performed by Novogene, Hong Kong.

#### Library preparation and sequencing

RNA concentration was measured using Qubit^®^ RNA Assay Kit in Qubit^®^ 2.0 Flurometer (Life Technologies, CA, USA). Sequencing libraries were generated from a total amount of 3 μg RNA per sample using NEBNext^®^ Ultra^™^ RNA Library Prep Kit for Illumina^®^ (NEB, USA) following manufacturer’s recommendations. The clustering of the index-coded samples was performed on a cBot Cluster Generation System using HiSeq PE Cluster Kit cBot-HS (Illumina) according to the manufacturer’s instructions. After cluster generation, the library preparations were sequenced on an Illumina HiSeq platform and 125 bp/150 bp paired-end reads were generated.

#### Quality control

Raw FASTQ reads were first processed through in-house Perl scripts. Clean reads were obtained by removing reads containing adapter, reads containing poly-N and low quality reads. To ascertain the quality of the reads, the Q20, Q30 and GC content were calculated. All downstream analyses were based on the clean data with high quality.

#### Reads mapping

To map the reads to their respective genes, the Ensembl reference genome release 79 for Rattus Norvegicus was used^68^. Index of the reference genome was built using Bowtie v2.2.3 and paired-end reads were aligned to the reference genome using TopHat v2.0.12. HTSeq v0.6.1 was used to count the number of reads mapped to each gene; the quantification of gene expression (Supplementary File 1).

### RNAseq data exploration

The read count file generated by Novogene was used as input for further analyses using DESeq2 v1.22.2, Bioconductor release 3.8^69^ in R v3.5. First, the data was pre-processed with a variance stabilizing transformation and explored using heat maps of the highest expressing genes and the sample-to-sample distances, and through principal component analysis (PCA). The PCA plot showed that samples from the medial prefrontal cortex cluster separately from the ones from the basolateral amygdala. However, there was one BLA sample that did not cluster with any other samples, suggesting its overall gene expression pattern is entirely different from all the other samples (Supplementary Figure 2A). Therefore, we excluded this sample from further analyses, leaving the cMV-Veh male BLA group with N=4.

### Gene Set Enrichment Analysis

To examine changes in sets of related genes, Gene Set Enrichment Analysis (GSEA) v4.0.1 was used through the desktop application from the Broad Institute^70^. The algorithm is developed to compare 2 groups or phenotypes, and is not suitable for analysis of interactions like we did in the differential expression analysis. Therefore, to examine the effect of maternal adversity and FLX exposure on the expression of gene sets, we decided to run the GSEA on the control group (cMV-Veh) versus 1. the maternal adversity group (sMV-Veh) and 2. the FLX-exposed group (cMV-FLX) per combination of sex and brain area.

The algorithm requires a gene sets database, an expression dataset, and phenotype labels. The gene sets database was generated by generating a list of Gene Ontology (GO) terms for all genes in our gene expression data using the Biomart database system of Ensembl genes 95, dataset Rnor_6.0^68^. We included all 3 GO domains: cellular component, molecular function, and biological process. The expression dataset was a normalized gene count dataset that was pre-processed with a variance stabilizing transformation through DESeq2 for every combination of sex and brain area. The phenotype labels were simply the group identifiers (cMV-Veh vs sMV-Veh, or cMV-Veh vs cMV-FLX). The GSEA results yield a list of gene sets that differentiate between the 2 phenotypes, and also which genes within these sets contribute the most to the observed difference. The Normalized Enrichment Score (NES) is the degree to which the gene set is overrepresented in a phenotype. We set the cutoff for significance at False Discovery Rate (FDR) <0.25.

### Differential Expression Analysis

DESeq2 was used to identify the genes that are differentially expressed depending on maternal adversity exposure, FLX treatment, or a combination of both, a sMV + FLX + sMV*FLX model was applied to every combination of sex and brain area (PFC males, BLA males, PFC females, BLA females). Even though based on PCA, males and females do not differ in their global gene expression patterns (Supplementary Figure 2B), we decided to analyze them separately based on literature and previous findings from our lab^63^. The cutoff for statistical significance was set at FDR<0.1.

### DNA methylation analysis

To examine potential differential epigenetic regulation of genes of interest identified by differential expression analysis and GSEA (*Cldn11*, *Cnp*, *Mag* and *Mbp*), bisulfite pyrosequencing was performed on DNA originating from the same samples as were used for RNAseq. Bisulfite conversion of 240 ng genomic DNA was performed with EZ DNA methylation gold kit (Zymo Research, Leiden, The Netherlands) according to manufacturer’s protocol. This treatment converts cytosine but not 5-methylcytosine residues to uracil. Bisulfite-specific primers for the promoter regions of *Cldn11*, *Cnp*, *Mag* and *Mbp* were designed using PyroMark Assay Design Software 2.0 (Qiagen) (Supplementary Figure 3; Supplementary File 2). HotStarTaq master mix (Qiagen, Hilden, Germany) was used for amplification of 1 μL of the bisulfite-treated DNA modified using the following steps: DNA polymerase activation (95°C, 15 minutes), 3-step cycle of denaturation (94°C, 30 seconds), annealing (variable temperatures, 30 seconds), and extension (72°C, 30 seconds) repeated for variable number of cycles in a row (Supplementary File 2). The final extension was performed at 72°C for 7 minutes. The PCR product was analyzed for the extent of methylation per selected CpG positions by pyrosequencing using a PyroMark Q48 Autoprep Instrument (Qiagen). Finally, PyroMark Q48 software (Qiagen) was used to determine the methylation percentage of individual CpG positions. Of note, the DNA from 3 samples (2 cMV-Veh male, 1 cMV-FLX female) failed to be amplified during the PCR reaction. We therefore replaced these samples by others from the same treatment group but not previously used for RNAseq. CpG positions 2 and 7 for *Cnp* and CpG position 3 for *Mbp* had to be excluded because the measurements were unreliable (high peak height deviation). Since the percentage DNA methylation for the individual CpGs of the same gene correlated highly, we used the average percentage DNA methylation per gene for further analyses.

DNA methylation percentages for every combination of sex and brain area were visually inspected using a Q-Q plot in R v3.5 and deemed to be normally distributed. GraphPad prism v8 was used for linear regression analyses and two-way ANOVAs, with statistical significance set at *p*<0.05. Error bars in graphs represent SEM.

## Results

### Gene Set Enrichment Analysis reveals brain region-specific effects of perinatal fluoxetine and maternal adversity on myelin-related gene sets

First, we used Gene Set Enrichment Analysis (GSEA) to investigate the effects of maternal adversity and FLX exposure on the expression of gene sets related to a particular cellular, molecular or biological function. To this end, we compared the cMV-Veh group to the cMV-FLX group and to the sMV-Veh group for every combination of sex and brain area, yielding 8 comparisons (Supplementary File 3). For each comparison, we plotted the top 30 gene sets with the largest enrichment score (Figure 2A-D). In males, several key gene sets related to oligodendrocytes and myelin formation were positively associated with the cMV-FLX phenotype in the PFC, whereas they were negatively associated with the cMV-FLX phenotype in the BLA (Figure 2A). More specifically, FLX exposure was associated with an upregulation of the gene sets Oligodendrocyte differentiation (NES=2.38, FDR<0.001), Oligodendrocyte development (NES=2.25, FDR<0.05), Myelin sheath (NES=2.23, FDR<0.01) and Myelination (NES=2.12, FDR<0.05) in the PFC, but with a downregulation of the gene sets Myelin sheath (NES=−2.66, FDR<0.001), Oligodendrocyte differentiation (NES=−2.19, FDR<0.05) and Myelination (NES=−2.18, FDR<0.05) in the BLA (Figure 2A, Figure 3A).

**Figure 2:**
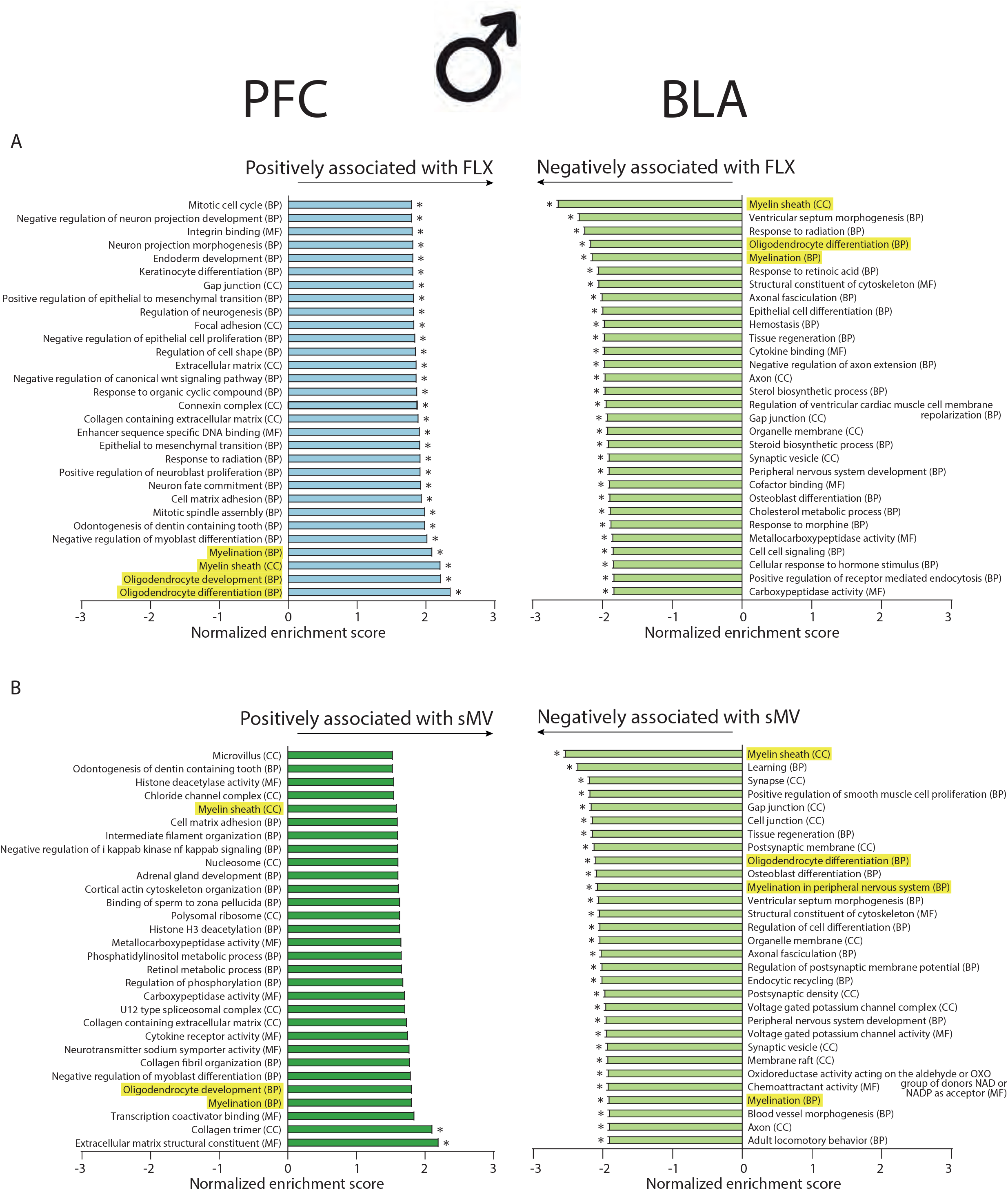
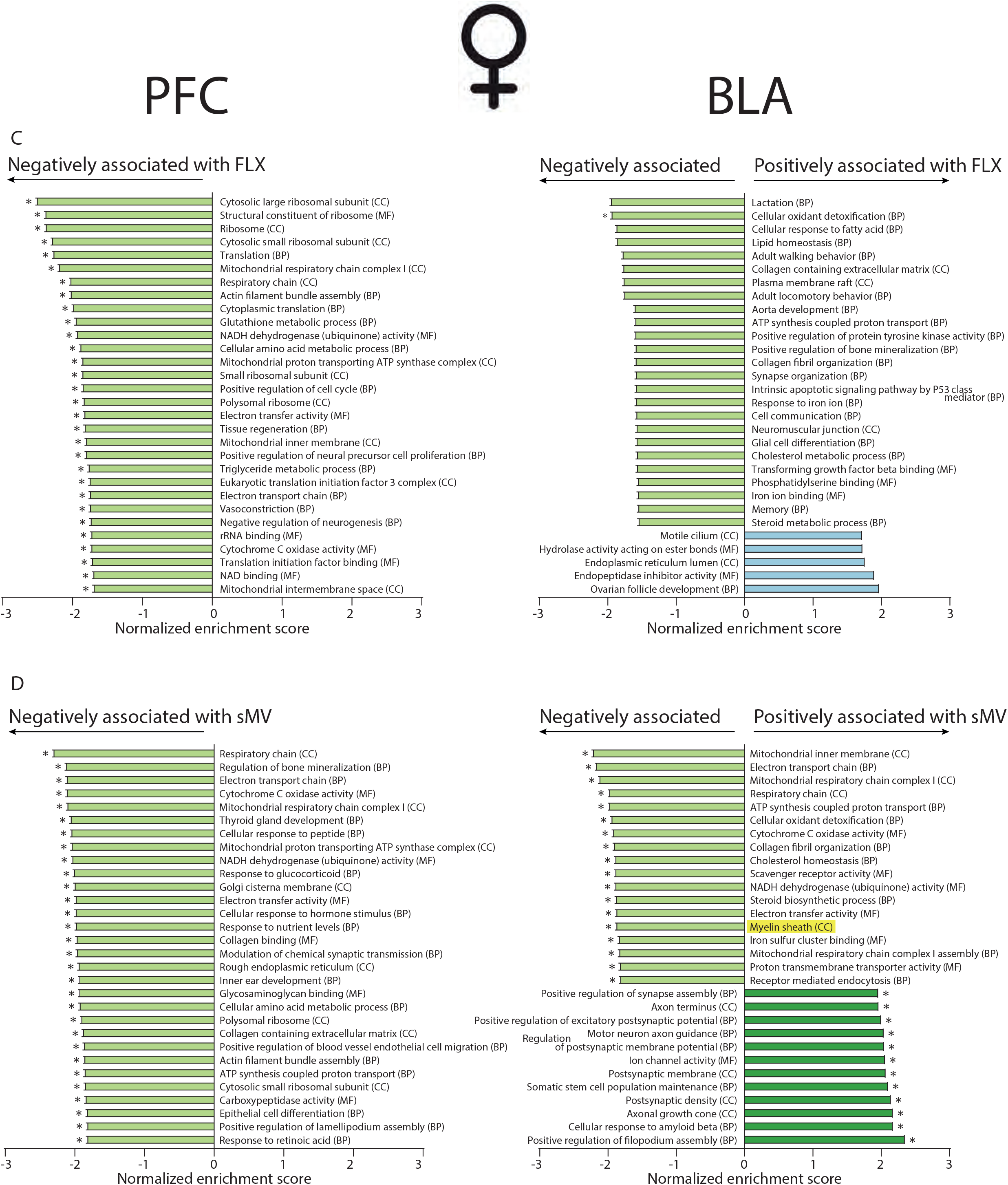
The top 30 gene sets associated with either of the 2 phenotypes being compared in the PFC and BLA. A.) The effect of fluoxetine exposure (cMV-FLX vs cMV-Veh) in males. B.) The effect of maternal adversity (sMV-Veh vs cMV-Veh) in males. C.) The effect of fluoxetine exposure (cMV-FLX vs cMV-Veh) in females. D.) The effect of maternal adversity (sMV-Veh vs cMV-Veh) in females. Myelin-related gene sets are highlighted in yellow. * indicates FDR<0.25. Related to Supplementary File 3.

**Figure 3:**
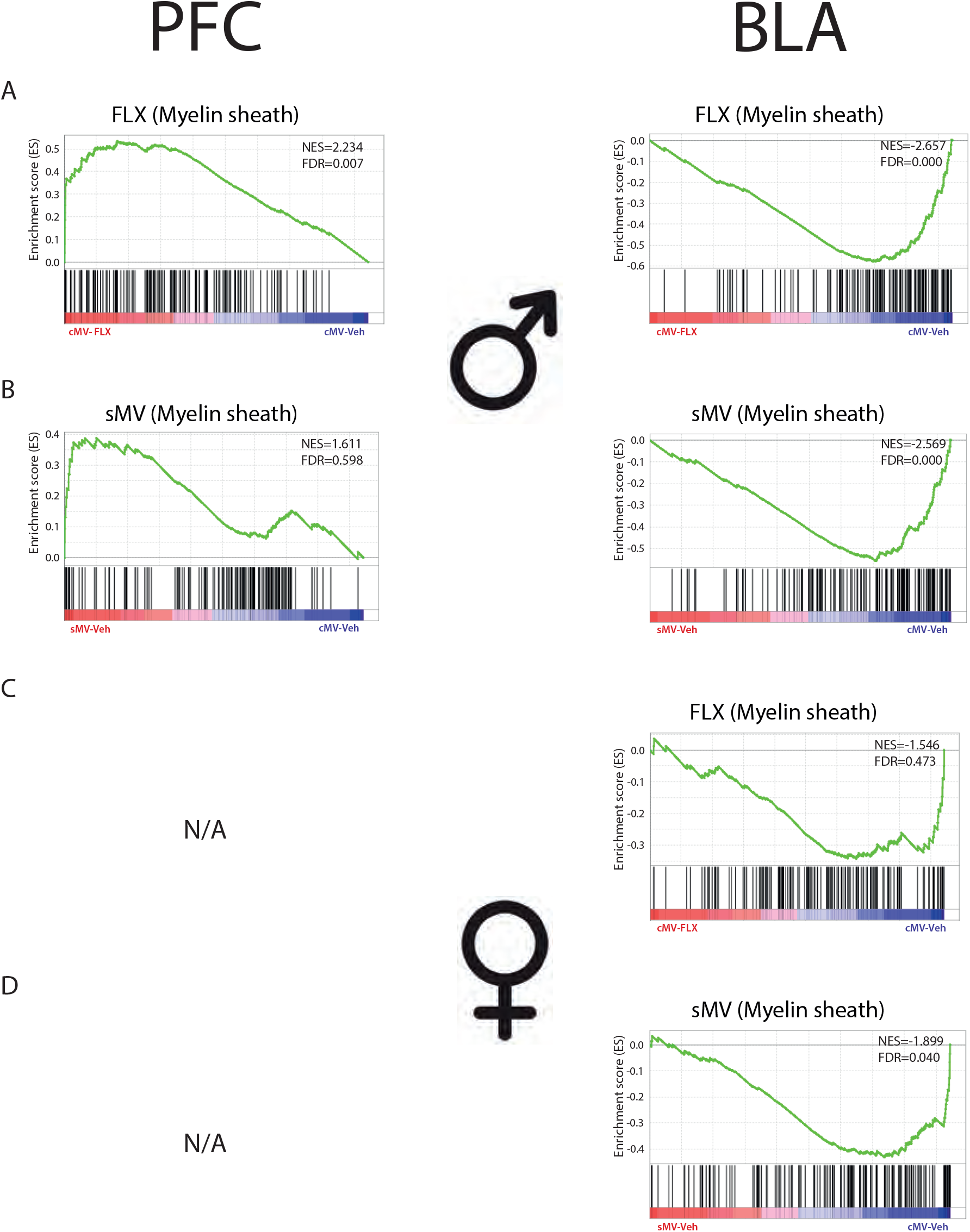
Enrichment plots for the gene set “Myelin sheath” in the PFC and BLA. The black vertical lines correspond to individual genes in this gene set, and their horizontal position corresponds to how strongly they associate with either of the two phenotypes that are being compared. A.) The effect of fluoxetine exposure (cMV-FLX vs cMV-Veh) in males. B.) The effect of maternal adversity (sMV-Veh vs cMV-Veh) in males. C.) The effect of fluoxetine exposure (cMV-FLX vs cMV-Veh) in females. D.) The effect of maternal adversity (sMV-Veh vs cMV-Veh) in females. NES = normalized enrichment score, FDR = false discovery rate. N/A indicates that the “Myelin sheath” gene set was not in the top 100 gene sets for this comparison. Related to Supplementary File 5.

Maternal adversity had a similar effect on gene set enrichment in males; myelin-related gene sets were among the top gene sets positively associated with sMV in the PFC, while they were negatively associated with sMV in the BLA (Figure 2B). In the PFC, maternal adversity was associated – although not with an FDR lower than 0.25 – with an upregulation of the gene sets Myelination (NES=1.83, FDR=0.47), Oligodendrocyte development (NES=1.83, FDR=0.39) and Myelin sheath (NES=1.61, FDR=0.60). In the BLA, however, maternal adversity was associated with a downregulation of the gene sets Myelin sheath (NES=−2.57, FDR<0.001), Oligodendrocyte differentiation (NES=−2.13, FDR<0.01), Myelination in peripheral nervous system (NES=−2.11, FDR<0.05) and Myelination (NES=−1.94, FDR<0.05) (Figure 2B, Figure 3B). Maternal adversity was also negatively associated with several gene sets related to neuronal communication in the male BLA, such as Learning (NES=−2.39, FDR<0.001) and Synapse (NES=−2.22, FDR<0.01) (Figure 2B).

In females, FLX exposure was negatively associated with an array of gene sets related to general cell maintenance and proliferation in the PFC, such as Ribosome (NES=−2.40, FDR<0.001) and Translation (NES=−2.29, FDR<0.001) (Figure 2C). In the BLA, there was only one significantly (FDR<0.25) downregulated pathway in the cMV-FLX females, which was Cellular oxidant detoxification (NES=−1.97, FDR<0.25) (Figure 2C). Although not significantly, the gene set Myelin sheath in the female BLA showed the same negative association with FLX as in the male BLA (NES= −1.55, FDR= 0.47) (Figure 3C).

Maternal adversity was associated with changes in many pathways in the female PFC. For example, gene sets related to endocrine signaling such as Thyroid gland development (NES=−2.08, FDR<0.05) and Response to glucocorticoid (NES=−2.03, FDR<0.05) were downregulated in the PFC in sMV females (Figure 2D). In the BLA, gene sets related to neuronal cell proliferation and synaptic plasticity were upregulated in sMV females, such as Postsynaptic density (NES=2.17, FDR<0.05) and Regulation of postsynaptic membrane potential (NES=2.06, FDR<0.05) (Figure 2D). In addition, like in the male sMV BLA, the gene set Myelin sheath was downregulated in the female sMV BLA (NES=−1.90, FDR<0.05) (Figure 2D, Figure 3D).

### Differential expression analysis reveals that maternal adversity and perinatal fluoxetine exposure interact to affect myelin-related gene expression in the BLA

Considering the similarity between the effects of maternal adversity and perinatal fluoxetine exposure on the enrichment of gene sets, we continued by examining potential interaction effects on the single-gene level. To this end, we constructed a FLX * maternal adversity interaction model using DESeq2 for every combination of sex and brain area, yielding 4 models (Supplementary File 4). Overall, FLX was associated with the most significantly differentially expressed genes in the male BLA (17 genes at FDR<0.1, Supplementary Figure 4B), and maternal adversity was associated with the most significantly differentially expressed genes in the female PFC (29 genes at FDR<0.1, Supplementary Figure 4C). In the BLA of both males and females, the top genes that showed a FLX * maternal adversity interaction effect were related to myelin, although not significantly (FDR<0.1) in females (Supplementary File 4). Some of these genes were also the main contributors to the effects found on the gene set Myelin sheath (Supplementary File 5, Figure 4). Specifically, claudin-11 (*Cldn11*), 2’,3’-cyclic-nucleotide 3’-phosphodiesterase (*Cnp*), myelin−associated glycoprotein (*Mag*) and myelin basic protein (*Mbp*) showed a FLX * maternal adversity interaction effect in the BLA (Figure 4B,4D). All of these genes were downregulated by FLX in cMV animals, but upregulated by FLX in sMV animals in the BLA. In the PFC, no significant effects of FLX and maternal adversity on the expression of *Cldn11*, *Cnp*, *Mag* and *Mbp* were found (Supplementary File 4, Figure 4A,4C).

**Figure 4:**
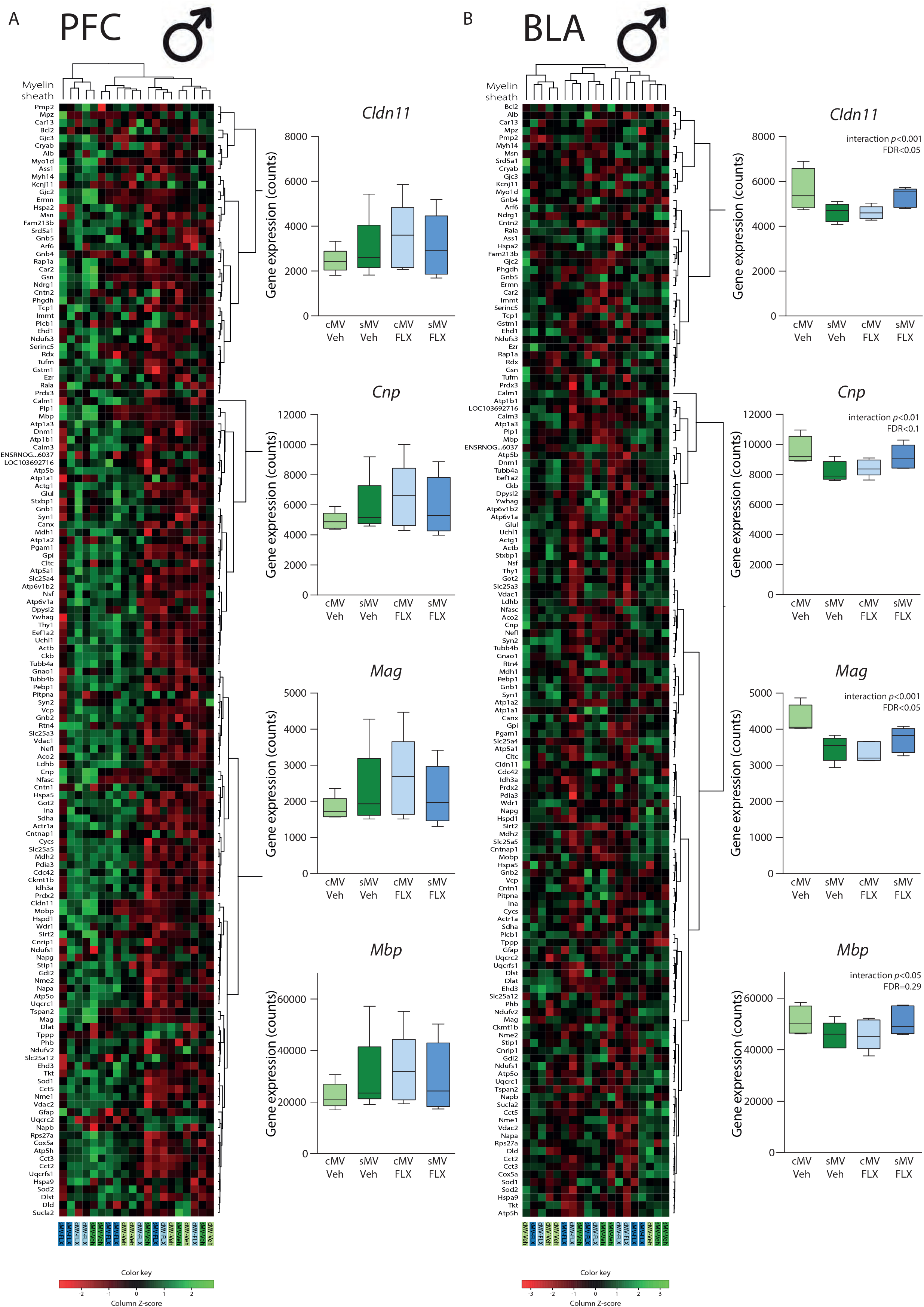
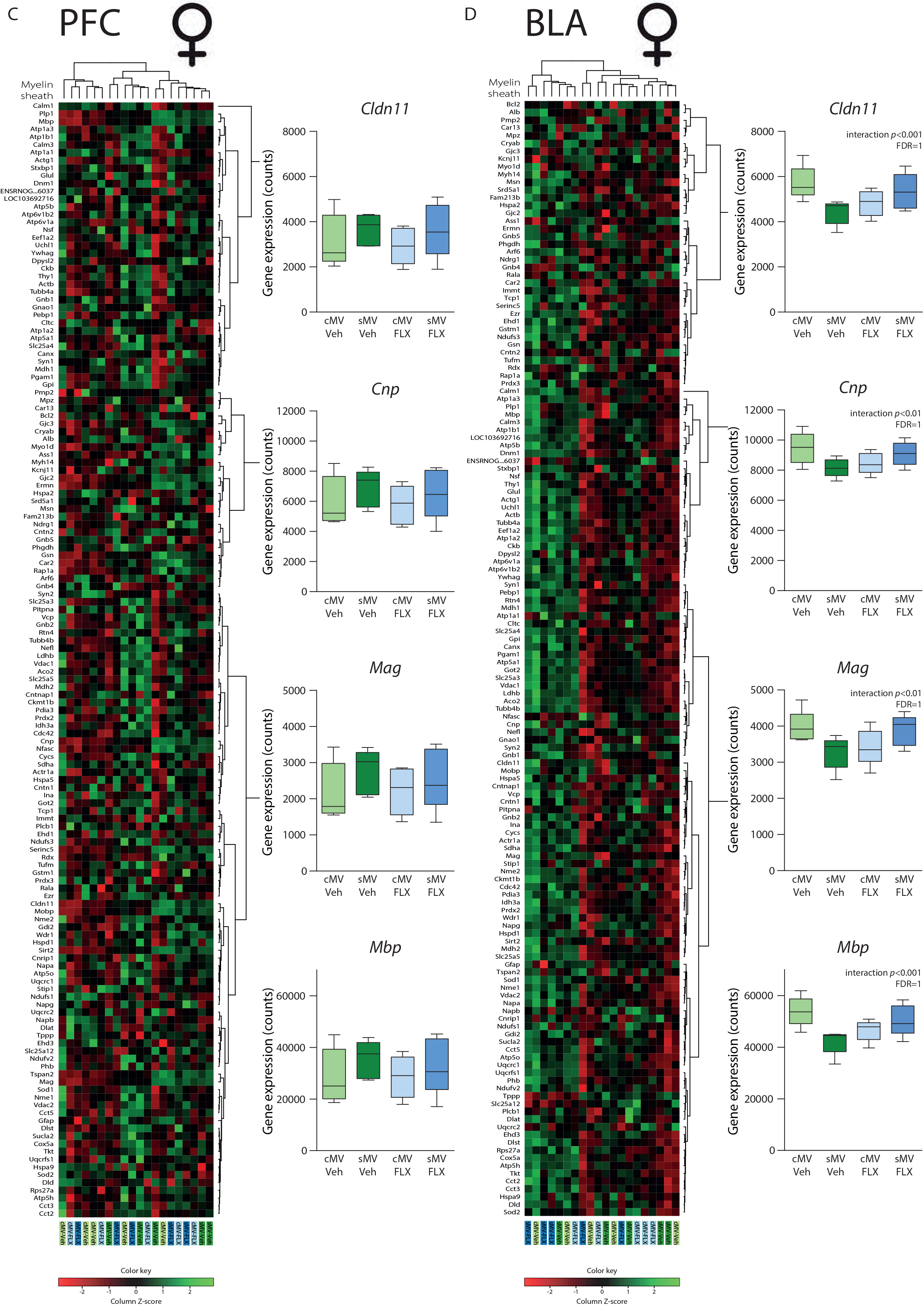
Heat maps of the expression of all individual genes in the gene set “Myelin sheath”, and bar plots of the gene counts of 4 selected genes: *Cldn11*, *Cnp*, *Mag* and *Mbp* for: A.) The male PFC. B.) The male BLA. C.) The female PFC. D.) The female BLA. Related to Supplementary File 1, 4, and 5.

### DNA methylation of *Mag* and *Mbp* correlates to gene expression

To examine epigenetic regulation potentially underlying the effects of FLX and maternal adversity on myelin-related gene expression, we next examined DNA methylation levels around the transcription start sites of *Cldn11*, *Cnp*, *Mag* and *Mbp* (Supplementary Figure 3, Supplementary File 6). We used DNA samples from the same tissue punches we also used for RNA sequencing, allowing for examination of correlations between DNA methylation and expression levels of these genes. Linear regression analyses showed there was a significant negative correlation between DNA methylation and *Mag* gene expression in both the PFC (R^2^=0.1689, *p*<0.05, Figure 5A) and the BLA (R^2^=0.2299, *p*<0.01, Figure 5B). There were no significant interaction effects of FLX * maternal adversity on *Mag* DNA methylation percentages, but a strong trend in the female PFC (*p*=0.059) (Figure 5A,5B). For *Mbp*, there was a negative correlation between DNA methylation and gene expression in the PFC (R^2^=0.2525, *p*<0.01, Figure 5C) but not in the BLA (R^2^=0.0000, *p*=0.9597, Figure 5D). There was a significant main effect of FLX on PFC Mbp methylation percentages in females (Figure 5C,5D). Unlike for *Mag* and *Mbp*, there were no correlations between DNA methylation and gene expression of *Cldn11* and *Cnp* (Supplementary Figure 5), so these were not further examined for FLX * maternal adversity interactions.

**Figure 5:**
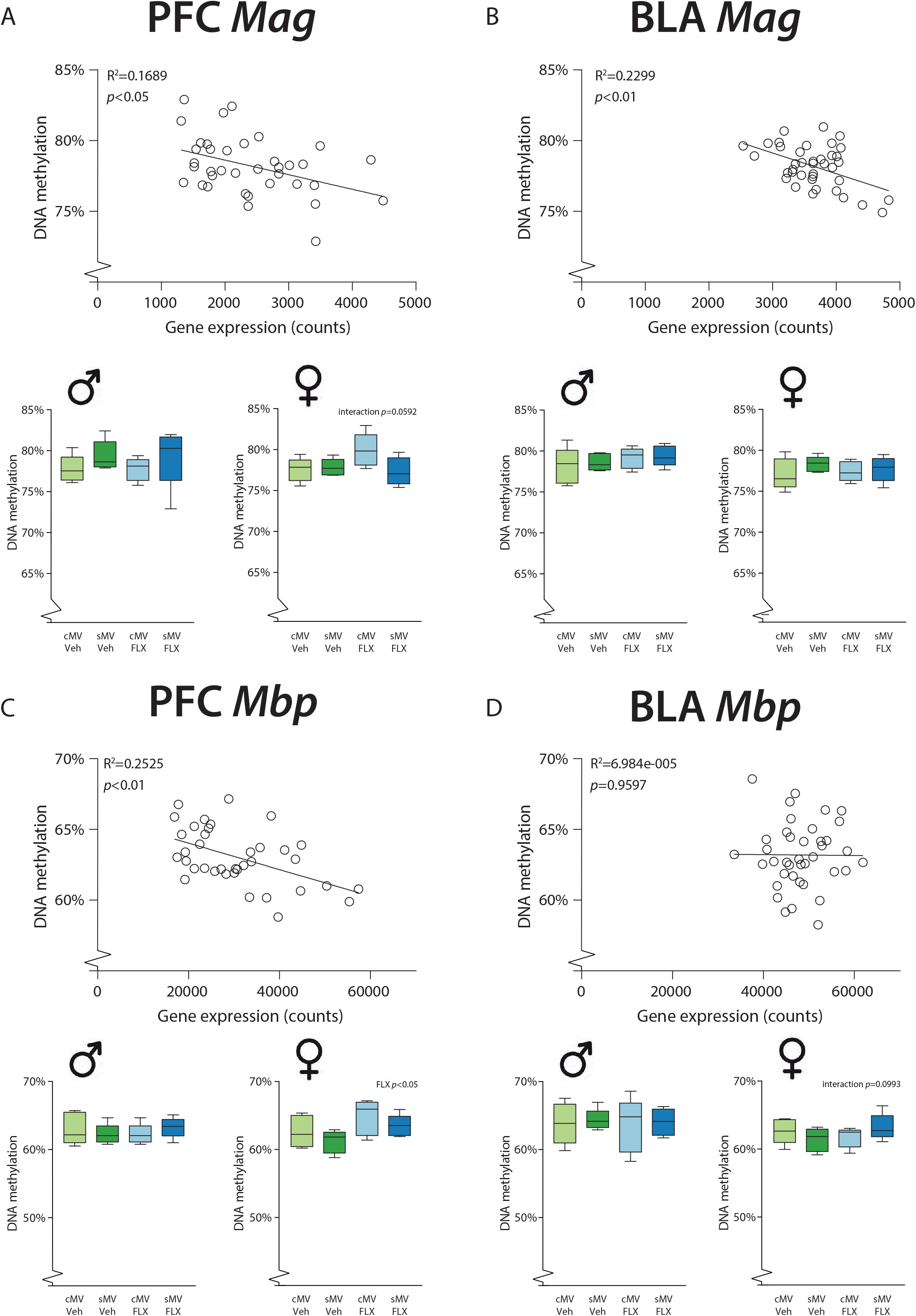
Scatter plots of DNA methylation percentages and gene counts, and bar plots of DNA methylation percentages in males and females for: A.) *Mag* in the PFC. B.) *Mag* in the BLA. C.) *Mbp* in the PFC. D.) *Mbp* in the BLA. Related to Supplementary File 1 and 6.

## Discussion

We investigated transcriptomic alterations in the prefrontal cortex (PFC) and basolateral amygdala (BLA) in male and female juvenile rats after exposure to perinatal fluoxetine and/or maternal adversity. In these key regions in the corticolimbic system of the brain, our interventions altered few genes significantly at the individual level. However, we found evidence for subtle yet consistent alterations in larger sets of genes related to myelination. Specifically, in males, myelin-related gene expression was upregulated in the PFC after perinatal fluoxetine exposure, while it was downregulated in the BLA. Myelin-related genes were not affected in the female PFC. The female BLA appeared to show a similar response to perinatal fluoxetine as the male BLA but it was weaker and non-significant. Interestingly, maternal adversity affected myelin-related gene expression in a manner similar to perinatal fluoxetine exposure. Differential expression analysis showed that maternal adversity and perinatal fluoxetine exposure interact to affect expression of genes such as myelin−associated glycoprotein (*Mag*), myelin basic protein (*Mbp*), claudin-11 (*Cldn11*) and 2’,3’-cyclic-nucleotide 3’-phosphodiesterase (*Cnp*). Gene expression correlated with DNA methylation of *Mag* and *Mbp*, highlighting epigenetic regulation as a potential mediator of perinatal fluoxetine-induced alterations in myelination.

Myelination of axons by oligodendrocytes plays a critical role in brain development and functioning; it allows for fast nerve conduction, reduces energy consumption, influences neuronal circuit formation and provides metabolic support to neurons^71^. In humans, myelination starts in mid-to-late gestation, with the majority of myelination taking place in the first two years of life^72^. In rodents, myelination starts after birth and spreads throughout the brain in a coordinated fashion during the second postnatal week^73^. At PND21, the limbic system is among the last regions to become fully myelinated, a process which takes another few weeks^73^. Efficient myelination is a complex process, requiring the interplay of numerous molecules, such as MAG, MBP, CLDN11 and CNP, to form and maintain multiple lipid layers around the axon^71^. MAG attaches the myelin sheath to the axon. Meanwhile, MBP binds the membrane layers of myelin to aid in its compacted structure. CLDN11 composes the tight junctions connecting the outer layers, thereby playing a role in the barrier function of myelin. To facilitate diffusion of molecules, CNP counteracts MBP to create cytoplasmic channels within the myelin sheath. Our results suggest that developmental exposure to SSRIs and maternal adversity interfere with myelination at a critical time in development, although long-term consequences of these alterations remain to be investigated.

The observation that perinatal fluoxetine exposure affects myelin-related gene expression is also evident by findings of Kroeze et al. who treated female rats with fluoxetine from PND1 to PND21. They found that myelin-related genes ciliary neurotrophic factor (*Cntf*) and transferrin (*Tf*) were downregulated in the adult hippocampus^74^. The same group characterized developmental transcriptomics in the medial PFC in male SERT knockout rats, mimicking SERT inhibition by SSRIs^75^. Constitutive SERT knockout decreased *Cldn11*, *Mag* and *Mbp* expression at PND8, increased it at PND14, and decreased it again at PND21 versus wildtypes. This is contrary to our findings of enhanced PFC myelin-related gene expression after fluoxetine exposure at PND21, despite the punched area being identical, highlighting that total lack of the SERT is not equal to SERT inhibition with SSRIs during the early development. Simpson et al. treated rats with the SSRI citalopram in the perinatal period and detected both hypo- and hypermethylation in the corpus callosum in adulthood^46^. In addition, they observed higher levels of abnormal axons after developmental SSRI exposure, with more severe deficits in males than in females^46^. Work in SERT knockout versus wildtype rats also indicated lower connectivity in the corpus callosum, while white matter integrity overall was mostly intact^76^. Overall, the effect of developmental SSRI exposure or constitutive SERT knockout on myelination appears to be highly dependent on brain region, age and sex.

It might be that a direct pathway links SSRI exposure to alterations in myelination. Consistent with this idea, *in vitro* evidence showed that increased serotonin levels damage immature oligodendrocytes^77^. Alternatively, high serotonin levels might affect myelination indirectly by affecting axons^77^. The onset of myelination is linked to neuronal differentiation^78^ and its unfolding controlled by neural acitivity^72^. Moreover, alterations in serotonin availability during neurodevelopment change the maturation of thalamocortical axons^79^, the medial PFC^44^, and the PFC to dorsal raphe nucleus circuit^45^. Increased perinatal serotonin levels are associated with lower levels of reelin, a glycoprotein involved in neuronal migration and cortical organization, in the rat brain^80^ and human umbilical cord^81^. It has been suggested that, considering reelin levels usually decrease across development, this may indicate an accelerated rate of neurodevelopment^81^. In line with this notion, developmental citalopram treatment is associated with earlier onset of synaptogenesis^82^. In fact, *in vitro* evidence suggests that embryo development is accelerated in human embryos treated with fluoxetine^83^. Similarly, the brain region- and developmental stage-specific effects of perinatal SSRI exposure might be signs of accelerated brain maturation. We speculate that both the BLA and PFC myelinate earlier due to SSRI exposure, with the peak of BLA myelination shifting to before PND21 (and thus leading to a *decrease* in myelination at this time), and PFC myelination shifting to around PND21 (leading to an *increase* in myelination at this time).

Further support of a fairly unspecific effect of fluoxetine on the rate of brain maturation is provided by our finding that the effects of maternal adversity are remarkably similar to those of perinatal fluoxetine exposure. This was true especially in the BLA, where myelin-related genes were also downregulated in the groups exposed to maternal adversity relative to controls. In line with this, the developing amygdala in both humans and animals is vulnerable to the effects of prenatal stress on functional and structural connectivity^84^. Although we did not find significant effects of maternal adversity on myelin-related gene expression in the medial PFC, other studies have identified an acute *increase* in myelination-related genes including *Mag*, *Mbp*, *Cldn11* and *Cnp* after early postnatal stress at PND15^85^, but a *decrease* in these same myelin-related genes and their proteins in adulthood^86^. This again suggests an effect on brain maturation, evidenced by a precocious differentiation of oligodendrocytes right after early life adversity, associated with a depletion of oligodendrocyte precursors in adulthood^85^.

Our lab previously identified interactions between maternal adversity and fluoxetine on a behavioral level^63^. In the current study, maternal adversity and fluoxetine interacted to affect *Mag*, *Mbp*, *Cldn11* and *Cnp* gene expression, especially in the BLA. The group receiving both treatments resembled the control group, suggesting that fluoxetine might normalize the effect in offspring exposed to *in utero* maternal adversity, at least on the expression of these myelin-related genes. Although the mechanism behind this is not clear, other work suggests that perinatal SSRI exposure is indeed capable of preventing some of the adverse effects of maternal adversity. For example, *in utero* citalopram exposure normalized the increased fetal frontal cortex serotonin levels induced by chronic prenatal stress^53^. Similarly, perinatal fluoxetine exposure largely prevented the effects of pre-gestational maternal stress on increased serotonin levels in the PFC in pre-adolescent offspring^49^, and stress coping behavior and hippocampal neurogenesis in adolescence^54^.

Preclinical evidence in rodents suggests that males are more vulnerable to develop long-term behavioral effects after perinatal SSRI exposure than females^87^. For example, our lab has identified stronger effects of perinatal fluoxetine exposure on social behavior in males than in females^63^. The current results also point to a stronger phenotype in males. In females, the dampening effect of fluoxetine on myelin-related genes in the BLA is present but non-significant, whereas myelin-related gene sets in the PFC are unaffected by fluoxetine. Connectivity studies in humans suggest that white matter development might occur earlier in girls than in boys^72^, potentially modulating the ability of SSRIs to alter this process. Whether the male bias seen in behavioral outcomes of early SSRI exposure is related to altered myelination during development remains to be determined. Prenatal stress has been associated with sex-specific neurodevelopmental outcomes as well^49,52^. Our results suggest that the transcriptomic state of the male PFC at PND21 is rather insensitive to the effects of maternal adversity, while a substantial number of genes and gene sets are differentially regulated in this area in female offspring exposed to maternal adversity. It remains to be seen whether this female bias in rats is related to the stronger effect in girls compared to boys found in human neuroimaging research after *in utero* exposure to maternal depressive symptoms^31–33^.

Epigenetic regulation of gene expression, in particular by DNA methylation at CpG dinucleotides, is thought to be a mediator between early life experiences and later-life health and behavior^88^. Because measuring DNA methylation in the human brain is only possible when postmortem tissue is available^89^, it is usually measured in blood, placental tissue, or buccal samples^90^. Whole-genome DNA methylation analyses in such samples did not identify effects of SSRI exposure^91^, and effects of maternal depression on offspring DNA methylation are difficult to replicate^92^. Instead, we investigated the brain directly and took a candidate gene approach. We found negative correlations between gene expression and DNA methylation for *Mag* in the PFC and BLA, and for *Mbp* in the PFC. No such correlations were found for *Cldn11* or *Cnp*, potentially because the overall DNA methylation levels around these promoters were low. Others have suggested that DNA methylation contributes to the differentiation of oligodendrocyte precursor cells to mature oligodendrocytes^93^. Accordingly, mice with a knockout of DNA methyl transferase 1 in the oligodendrocyte lineage showed widespread hypomyelination. In agreement with the current results, they found a negative correlation between *Mag* expression and DNA methylation in oligodendroglial-lineage cells^93^. Overall, we provide the first indication that fluoxetine-induced changes in myelination might be mediated by an epigenetic mechanism.

The current study has several key strengths. First, we investigated both the PFC and BLA, which revealed a clear brain region-dependent effect on myelin-related gene expression after exposure to perinatal fluoxetine exposure and maternal adversity. Second, we studied both sexes, whereas this is still surprisingly uncommon in animal research^87^ as well as human imaging studies^84^, despite strong indications for sex-specificity. Third, we used the same tissue punches for transcriptomic analyses and targeted DNA methylation analyses, enabling direct correlations. Besides these important factors our study also has limitations to be addressed. First, we only assessed transcriptomics at one point in time, 24 hours after treatment cessation. We were specifically interested in PND21, because it is known that corticolimbic circuit development is sensitive to the environment in early postnatal life in humans, and in the analogous period in rodents^94^. In addition, others have shown that the largest SSRI-induced effects on transcriptomics in the amygdala were found at PND21^61^. However, our approach precludes inferences about the long-term effects of perinatal SSRI exposure and/or maternal adversity. Second, in order to secure RNA quality, we did not isolate protein. Consequently, we were unable to validate results at the protein level.

Nevertheless, our results provide a potential molecular mechanism for findings of altered brain connectivity in humans after exposure to maternal SSRI use^25–27^ or depressive symptoms during gestation^29–31^. As outlined in the introduction, human studies often observe similar neurodevelopmental outcomes in children exposed to SSRIs as in children exposed to unmedicated depression, evoking questions about the source of the effects. Here, we observe striking similarities between the effects of SSRI exposure and maternal adversity, such as lower myelin-related gene expression in the BLA. The next step is to elucidate whether these similarities are mainly due to increased brain serotonin levels. Future studies are also necessary to delineate the time course of changes in myelination and white matter development after perinatal insults. Both the causes of altered myelination, such as differences in neuronal activity^85^, and its consequences, such as altered brain circuit formation^95^, deserve to be further explored. In addition, it is unclear whether and under what circumstances alterations in myelination might recover later in life^73^. Lastly, we advocate for continued cross-talk between animal and human studies. For example, imaging techniques in rodents can provide a link between (pre)clinical molecular studies and human neuroimaging findings^46,53^.

## Acknowledgments

We thank Judith Swart, Wanda Douwenga and Christa Reitzema-Klein for their assistance with the early life stress procedure and drug administration.

---

**Supplementary Figure 1:**
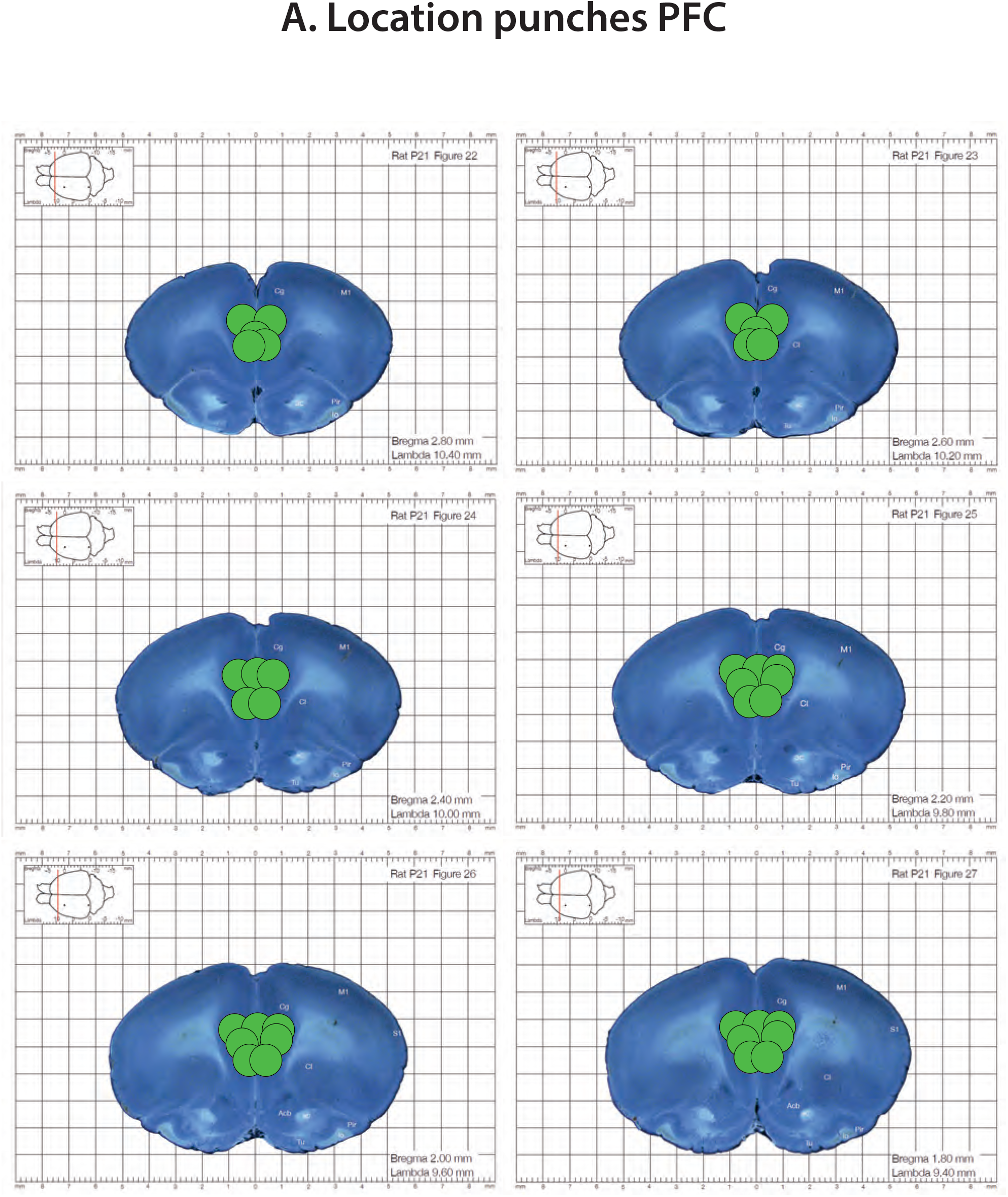
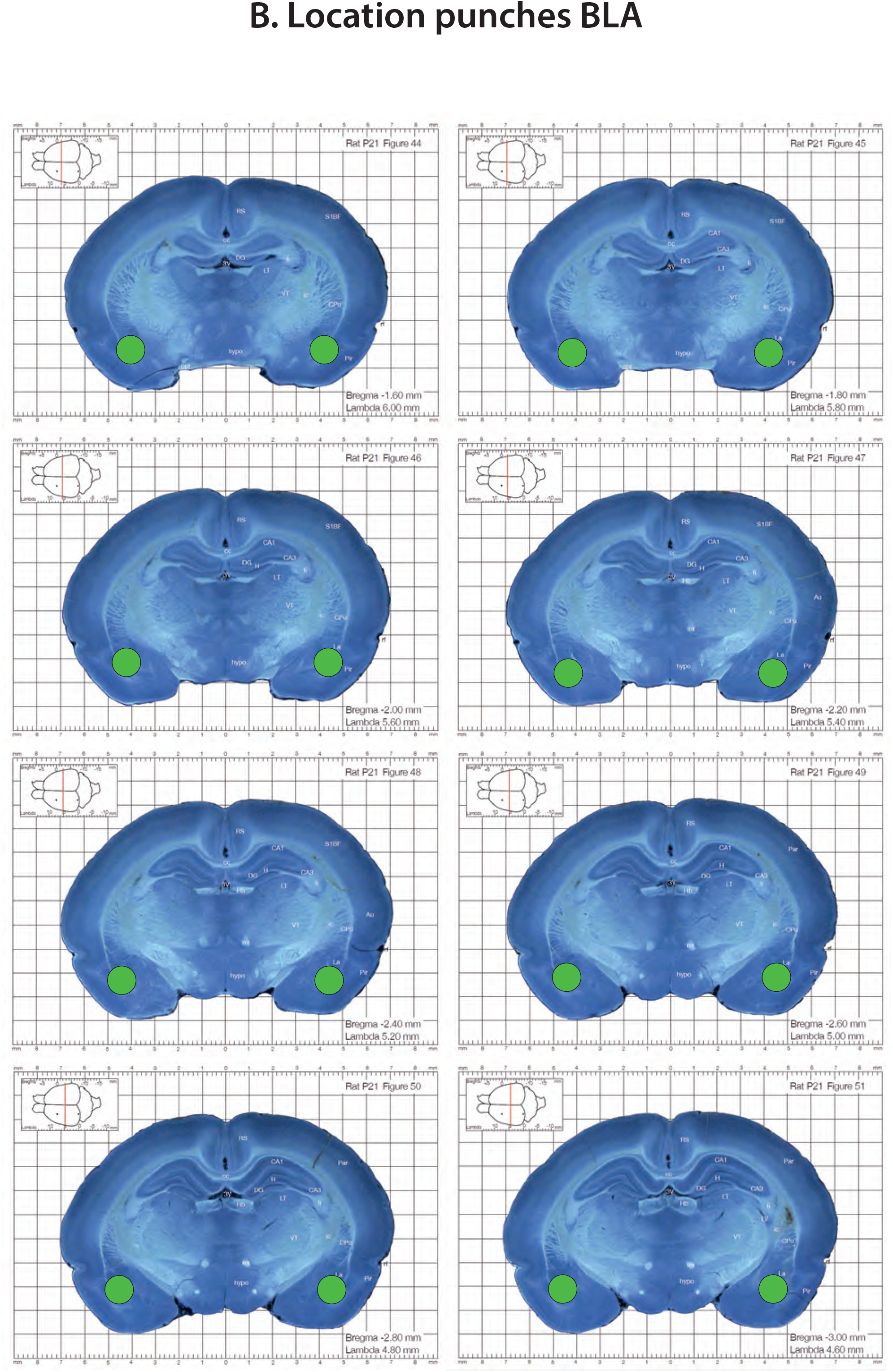
Location of the tissue punches, depicted in green in: A.) The PFC. B.) The BLA.

**Supplementary Figure 2:**
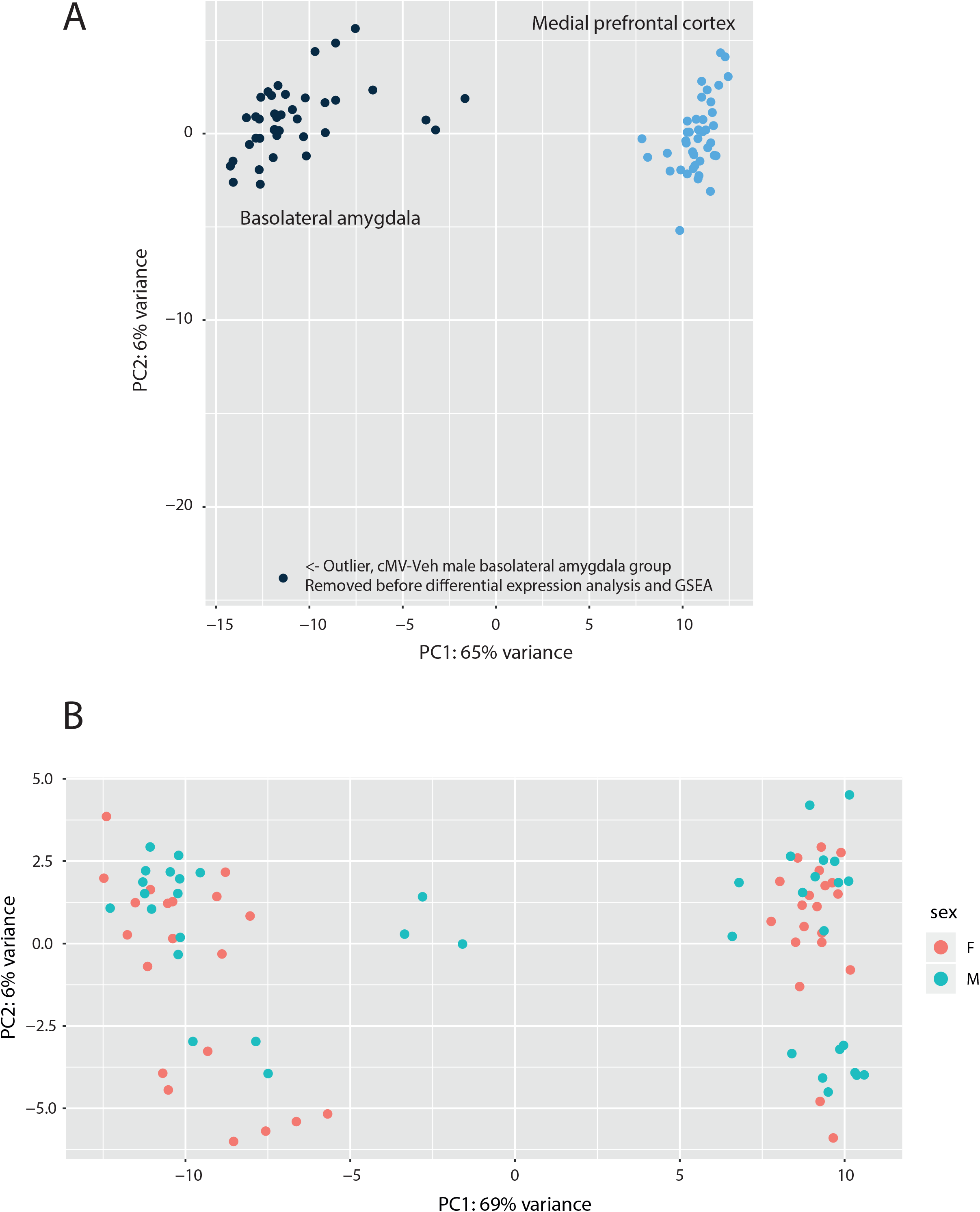
Principal coordinates (PC) plot of gene expression data, depicting: A.) The PFC and BLA cluster separately. One male cMV-Veh sample was determined to be an outlier and was removed before further analyses. B.) Males and females do not cluster separately. Related to Supplementary File 1.

**Supplementary Figure 3:**
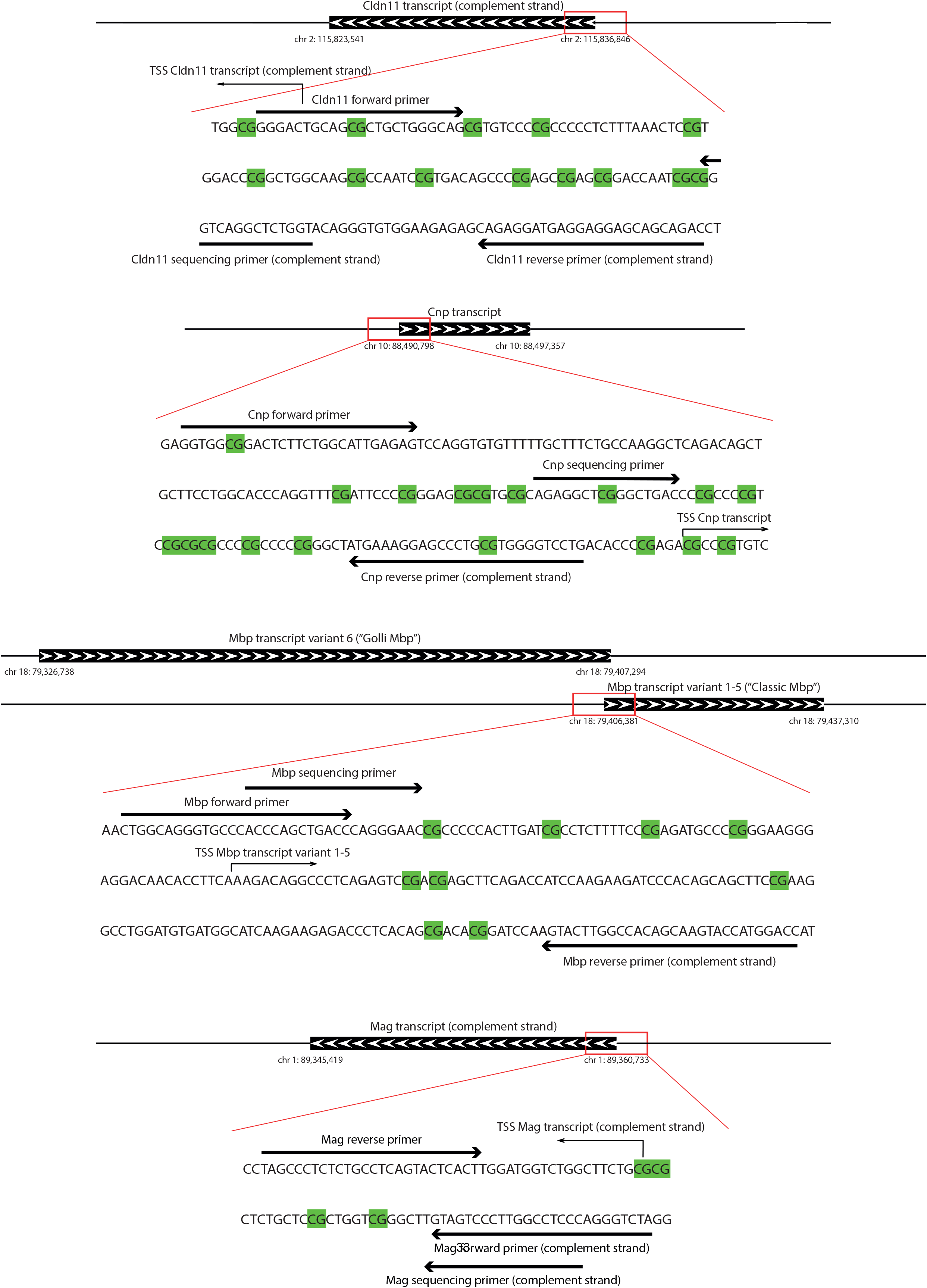
The genomic location of the primer targets used for DNA amplification and bisulfite sequencing. CpG locations are highlighted in green. A.) *Cldn11*. B.) *Cnp*. C. *Mag*. D. *Mbp*. Related to Supplementary File 2.

**Supplementary Figure 4:**
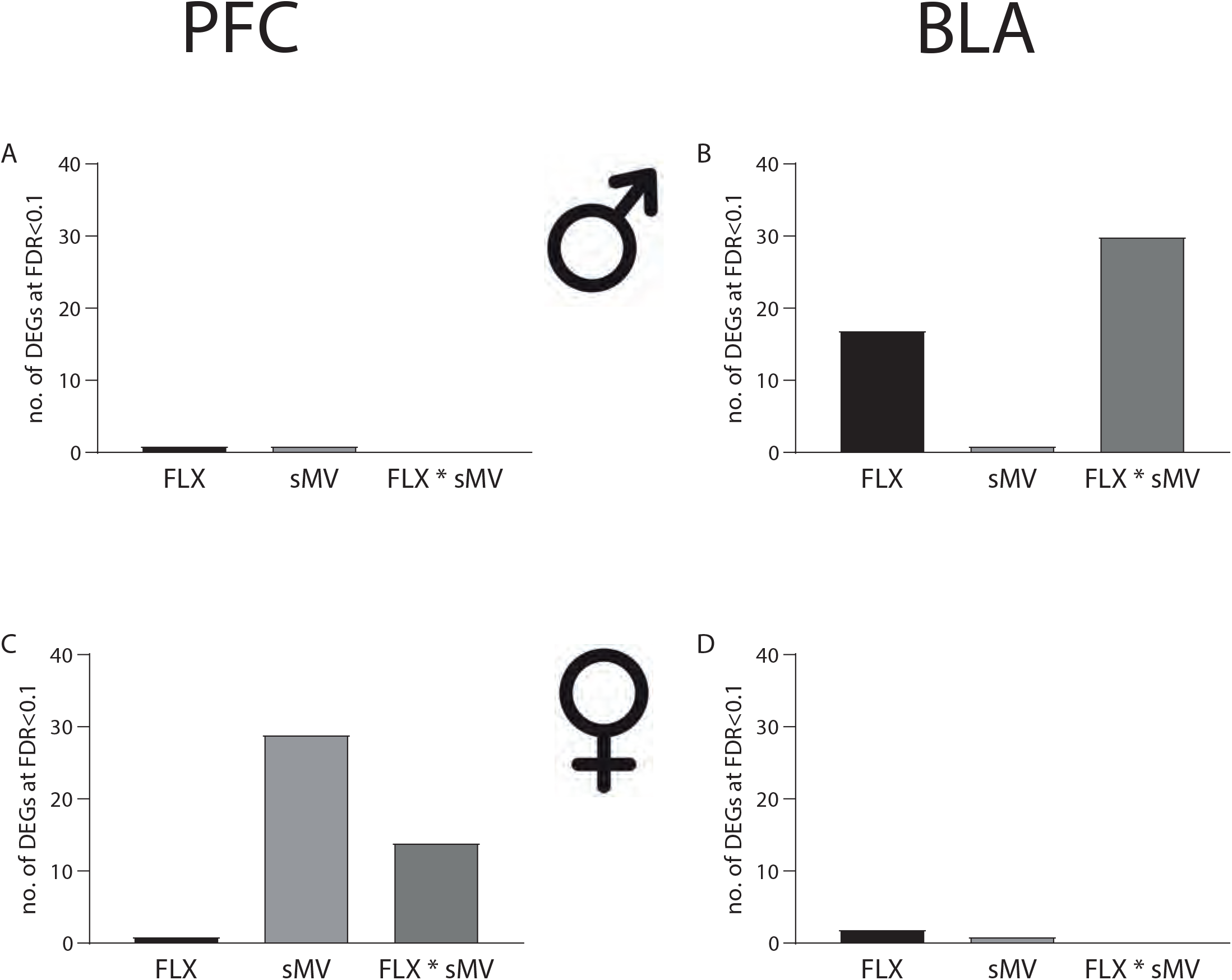
The number of differentially expressed genes (DEGs) at FDR<0.1 associated with FLX, sMV, or FLX*sMV, in: A.) The male PFC. B.) The male BLA. C.) The female PFC. D.) The female BLA. Related to Supplementary File 4.

**Supplementary Figure 5:**
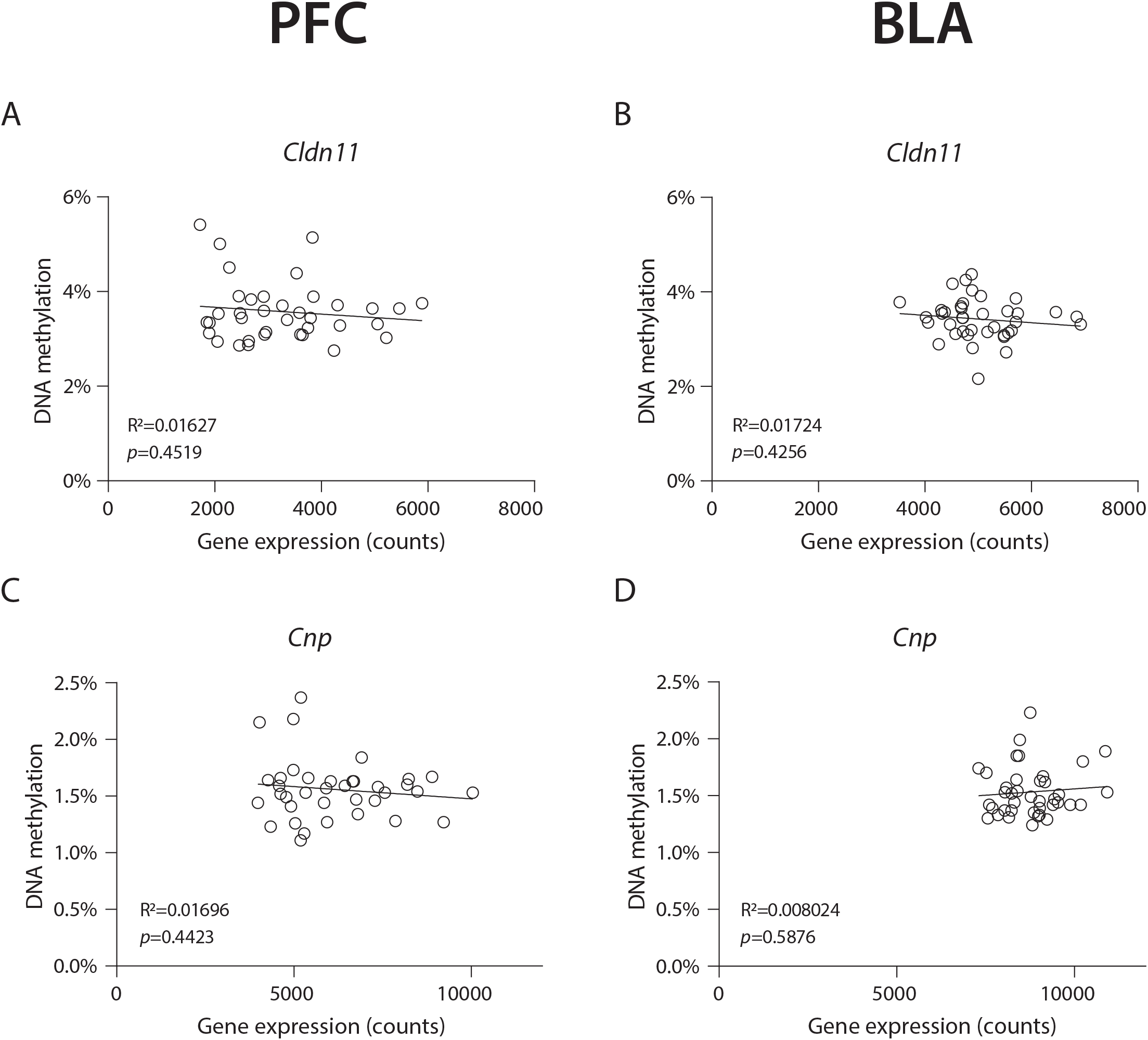
Scatter plots of DNA methylation percentages and gene counts for: A.) *Cldn11* in the PFC. B.) *Cldn11* in the BLA. C.) *Cnp* in the PFC. D.) *Cnp* in the BLA. Related to Supplementary File 1 and 6.

**Supplementary File 1:** Annotated read counts.

**Supplementary File 2:** Primer information.

**Supplementary File 3:** Gene Set Enrichment Analysis results.

**Supplementary File 4:** Differential expression analysis.

**Supplementary File 5:** Gene Set Enrichment Analysis results for Myelin sheath.

**Supplementary File 6:** DNA methylation results.

